# Functional organizational principle evaluated by microstimulation in the cortical facial motor areas

**DOI:** 10.1101/2025.03.04.641093

**Authors:** Sarina Karmacharya, Yifat Prut, Winrich A. Freiwald

**Affiliations:** Graduate School of Systemic Neurosciences, Ludwig-Maximilians-Universität München, Germany; Laboratory of Neural Systems, The Rockefeller University, New York, NY, USA; ELSC, The Hebrew University, Jerusalem, Israel

**Keywords:** Electrical microstimulation, cortical facial motor areas, facial expressions, facial movements, non-human primates

## Abstract

Facial gestures are crucial for social interactions among all primates. Since these actions are performed in the absence of direct visual feedback, their effectiveness relies on their stereotypical, almost reflexive patterns. One efficient mechanism for controlling ethological actions could involve motor cortical action maps (Graziano et al., 2002; 2016). Here, we studied whether stereotypical socio-communicative facial expressions are organized into discrete cortical zones by applying long-duration, supra-threshold microstimulation to fMRI-identified cortical areas of the facial motor system. Our findings revealed that these stimulation parameters produced complex facial responses by engaging multiple facial regions but did not elicit social-communicative facial behavior. The effects of long-duration stimulation appeared to be an amplified (both spatially and temporally) version of the responses seen with short-duration stimulation. The stimulation-evoked neural activity extended across the facial motor system, suggesting that these cortical regions function as part of a large-scale network designed to generate coherent, context-specific, and socially relevant motor outputs.

## Introduction

The somatotopic organization of the precentral gyrus is well-documented across many species, but its functional implications remain debated. This is likely due to contradictory findings from different studies that employed different experimental tools and protocols. Short-duration electrical stimulation triggered movements in specific body parts, reinforcing the idea of somatotopic representation (Fritsch and Hitzig, 1870; Penfield and Rasmussen, 1950). Another area of research proposes that the precentral gyrus includes distinct spatial zones that organize ethological actions when electrical stimulation is applied for durations matching the animal’s behavioral repertoire (Graziano et al., 2002; 2005; 2006; 2016). A recent fMRI study found that the motor cortex contains areas that represent both specific and multi-effector motor maps, potentially encoding sequences of behaviors frequently performed by the animal (Gordon et al., 2023). Thus, one efficient mechanism for controlling ethological actions could involve motor cortical action maps. Facial expressions are a form of stereotypical ethological actions, with the face serving as a key effector. These expressions are both prevalent and essential, particularly since primates use them to communicate reciprocally with species (Burrows, 2008; Waller et al., 2020). It follows then, that the facial cortical motor system (FCMS) can be a strong candidate for containing ethologically meaningful maps, and several studies have already identified ethologically relevant facial behaviors such as chewing, licking and defensive facial actions (Huang et al., 1989; Graziano et al., 2002; Cooke et al., 2003).

The core FCMS comprises five motor areas spanning the entire motor hierarchy, all of which project directly to the facial nucleus (Jenny and Saper, 1987; Welt and Abbs, 1990; Morecraft et al., 2001, 2004, 2007). Such a distributed network is suggestive that the production of facial expressions may depend on motor commands that evolve through hierarchical interactions rather than relying on localized, discrete action maps.

To determine the control scheme that is implemented in the FCMS, we implanted eight electrode arrays in cortical areas targeted by fMRI and systematically studied facial responses to suprathreshold long-duration stimulation. We found that long-duration stimulation in both low-level and high-level regions did not evoke ethologically meaningful communicative facial gestures. Our findings support the interpretation that the FCMS likely operates as a multi-area, hierarchically organized network.

## Results

To study the organizational principles of facial motor areas, we implanted eight chronic floating microelectrode arrays, guided by fMRI targeting, in two subjects (Fig. 1A). We assessed the causal contribution of these identified areas to facial behavior by observing facial responses to trains of electrical microstimulation delivered at two different durations (50 ms and 500 ms) and at suprathreshold current values (Fig. 1B). Behavioral responses were captured using videography and quantified by calculating optic flow, which measures the spatial extent of the evoked responses (Fig. 1C). Simultaneously, we recorded neural responses in all FCMS in response to concomitant stimulation delivered sequentially at each cortical site (Fig. 1D).

**Figure 1.**
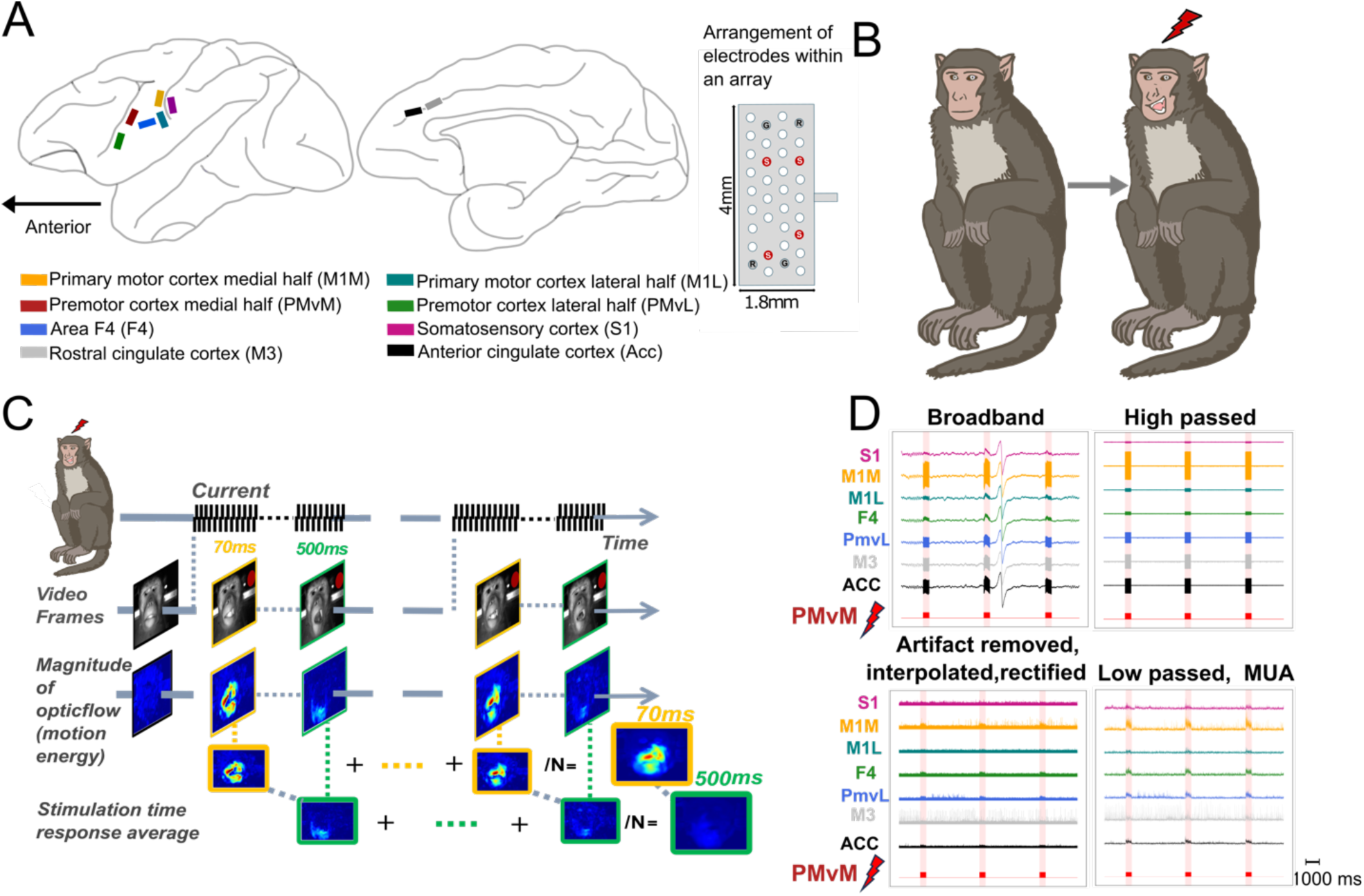
Experimental design. **A.** Schematic of recording and stimulation sites for brain of both subjects. Lateral and medial view of the right hemisphere are depicting the rough location for the implanted eight floating microelectrode arrays: one array in the somatosensory area, two arrays in the ventral portion of primary motor cortex, two arrays in the ventral portion of the premotor cortex and two arrays in the cingulate cortex. Each of the array had four stimulating electrodes labelled as “S” in red color, except for ACC which had two stimulating electrodes. **B.** Overview of the two experimental states observed in our paradigm to elicit behavioral responses. **C.** Optic flow magnitude was calculated from the temporally aligned video frames. Motion energy was averaged across trials to generate stimulation time response average map (STRA). **D.** Preprocessing of the simultaneously acquired neural signals in S1, M1M, M1L, PMvL, F4, M3 and ACC while stimulating a channel in PMvM. Time series were high pass filtered, stimulation related artifacts were removed, the missing signal was interpolated, rectified, and low passed filtered to get the final multi-unit activity (MUA) signal.

### Responses to long-duration trains evolve as a protraction of the responses obtained by short-duration trains

Past studies have suggested that sufficiently long trains of stimuli (i.e., durations comparable to natural movements) can reveal otherwise covert action maps. To investigate whether such action maps exist in the cortical motor system controlling the face, we compared facial responses evoked by short versus long stimulation trains. We constructed average response maps for 50 ms and 500 ms durations separately and normalized these maps (z-scored) using the mean and standard deviation calculated from the “without-stimulation” period at each pixel. Fig. 2A presents several examples of these maps.

**Figure 2.**
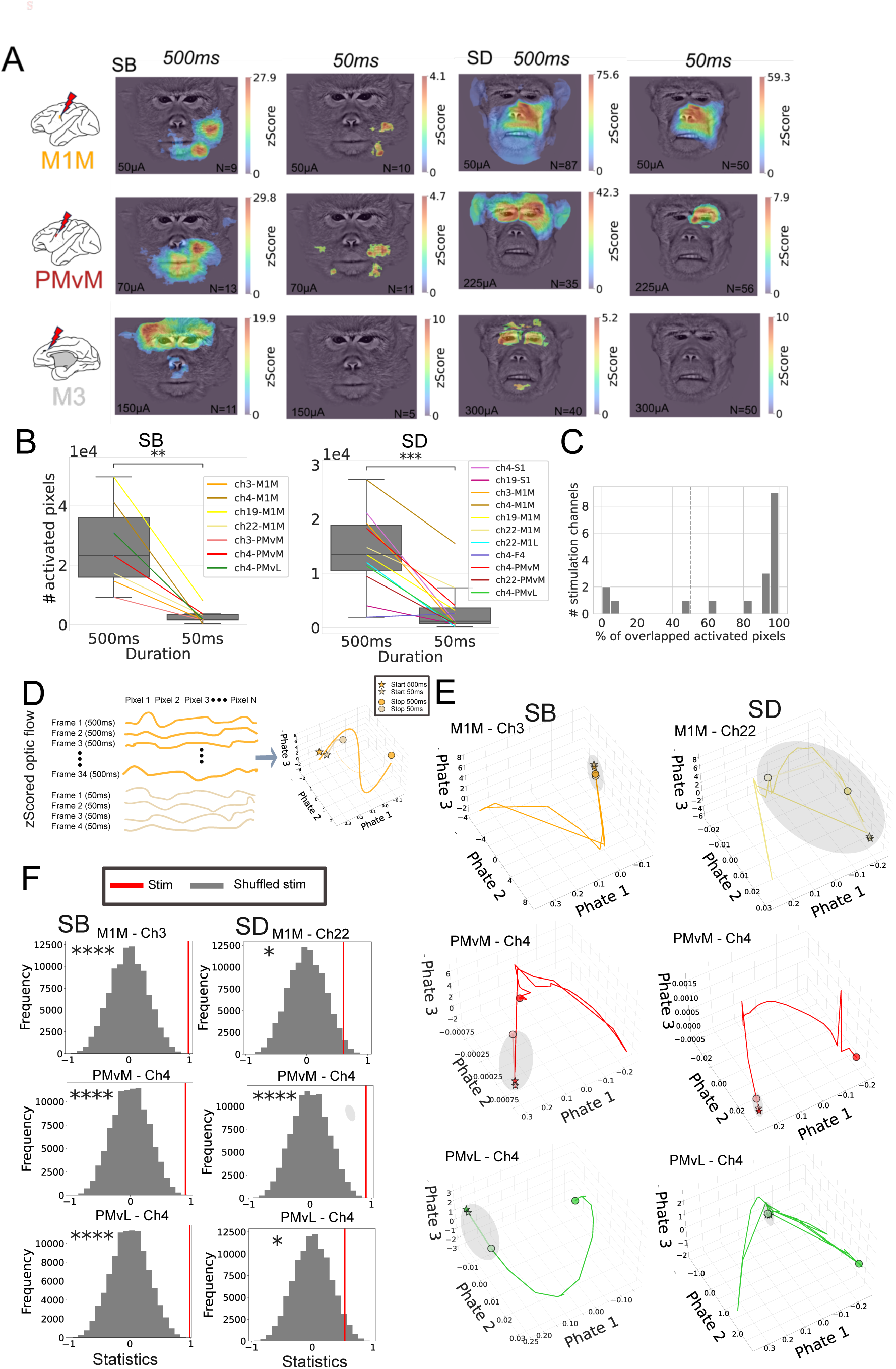
Stimulation evoked responses overlaid on the face. **A**. Evoked facial movements from three example sites from M1M, PMvM and M3. The STRA map was computed for each pixel within the face and z-scored by the data from “without-stimulation” epoch. The overlay heat map displays the maximum z-score across the entire stimulation period of 500 ms (Left panel) and 50 ms (Right panel) for each subject. The first two columns on the left show Monkey B and the last two columns show Monkey D. **B.** Distribution of significantly recruited pixels for 500 ms (Left) and 50 ms (Right) durations separately for each subject. The boxplot only displays channels that produced responses at both 50 ms and 500 ms, with line-connected data points indicating responses related to the same channel. Compared to 50 ms microstimulation, the long-duration condition (500 ms) significantly increased the number of recruited pixels (**p <0.01, ***p <0.001). **C.** Histogram of the overlapping pixels between 50 ms and 500 ms as a percentage of recruited pixels at 50 ms. **D.** Schematics of the Phate dimensionality reduction analysis on z-scored motion energy time-series for both (500 ms and 50 ms). **E.** Examples of Phate dimensionality reduction trajectories of experimental data for three stimulation channels. **F**. Permutation null distributions relative to the observed experimental statistic (red vertical line). The trajectories similarity across stimulation conditions was assessed with Spearman’s rank correlation. (Shuffled 100,000 times, *p<0.05, **p<0.01, ***p<0.001, ****p<0.0001), The left panel shows Monkey B and right panel shows Monkey D.

In these examples, short 50 ms stimulation trains (Fig. 2A, right panel) in medial M1 (M1M, upper row – Monkey B: 50 ms, 50 µA; Monkey D: 50 ms, 45 µA), medial PMv (PMvM, middle row – Monkey B: 50 ms, 70 µA; Monkey D: 50 ms, 225 µA), and M3 (lower row – Monkey B: 50 ms, 150 µA; Monkey D: 50 ms, 300 µA) led to spatially restricted facial responses or no visible response in both monkeys. Specifically, stimulation of M1M in both subjects evoked movements confined to the lower contralateral face. Stimulation of PMvM activated the contralateral orofacial region and a weaker activation of the ipsilateral side in Monkey B while eliciting responses in the contralateral eye regions in Monkey D. While stimulation of M3, no visible facial response was observed in either monkey during the stimulation duration.

In contrast to the responses observed at 50 ms, facial responses to long 500 ms (Fig. 2A, left panel) train durations exhibited an increased spatial activation pattern, as demonstrated in three examples: M1M (Monkey B: 500 ms, 50 µA; Monkey D: 500 ms, 50 µA), PMvM (Monkey B: 500 ms, 70 µA; Monkey D: 500 ms, 225 µA), and M3 (Monkey B: 500 ms, 150 µA; Monkey D: 500 ms, 300 µA). The number of activated pixels at 500 ms was significantly larger than at 50 ms in both monkeys (Fig. 2B, 2-tailed t-test, p < 0.01) for channels that exhibited activation at both durations. However, despite the increased activation with longer stimulation, there was considerable spatial overlap between the responses elicited by short and long trains (Fig. 2C).

To test the hypothesis that facial responses to short trains were essentially a truncated version of those obtained with long trains, we employed the Phate dimensionality reduction analysis (see Methods). This technique preserves both the global and local structures of data in a lower-dimensional space, which is particularly valuable for analyzing time-varying movement trajectories.

We began by computing the trial-average and z-score motion energy for both train durations (50 ms and 500 ms) at each stimulation site (Fig. 2D, left panel). We then calculated the Phate trajectories for both durations within the same space for each stimulation channel (Fig. 2D, right panel). Fig. 2E illustrates example trajectories representing the evoked movements for 50 ms and 500 ms stimulation durations. The gray ellipse is a visual aid to show the position of the 50 ms trajectory (Fig. 2E).

From these low-dimensional trajectories, we correlated the trajectories obtained from the 50 ms train with the first 50 ms segment of the trajectory from the long (500 ms) stimulation train. To determine the statistical probability of our observations, we conducted a permutation test by comparing the observed test statistics to the distribution of test statistics generated from permuting the data 100,000 times (Fig. 2F).

The examples shown in Fig. 2F (Left panel Monkey B and Right panel Monkey D) demonstrate a significantly high correlation between the trajectories obtained for the two durations for the example channels. The Spearman correlation coefficients and one-tailed permutation test results are as follows: Monkey B: M1M-ch3: rho = 0.98, p<0.0001; Monkey B: PMvM-ch4: rho = 0.91, p<0.0001; Monkey B: PMvL-ch4: rho = 0.98, p<0.0001; Monkey D: M1M-ch22: rho = 0.60, p<0.05; Monkey D: PMvM-ch4: rho = 0.91, p<0.0001; Monkey D: PMvL-ch4: rho = 0.53, p<0.05. These results are shown in Fig. 2F indicating that the evoked movements at 500ms duration represent a continuation of the movements associated with the 50ms stimulation.

### Long train electrical stimulation recruits multiple parts of the face and elicits complex motor responses

Next, we examined the complexity of responses evoked by long duration stimulation, defining complexity as the number of facial sub-regions recruited by the stimulation. We segmented the face into eleven regions and quantified the evoked responses in each sub-region (Fig. 3A). Fig. 3B shows examples of region-specific response patterns evoked by stimulation in a channel at PMvM for both subjects.

**Figure 3.**
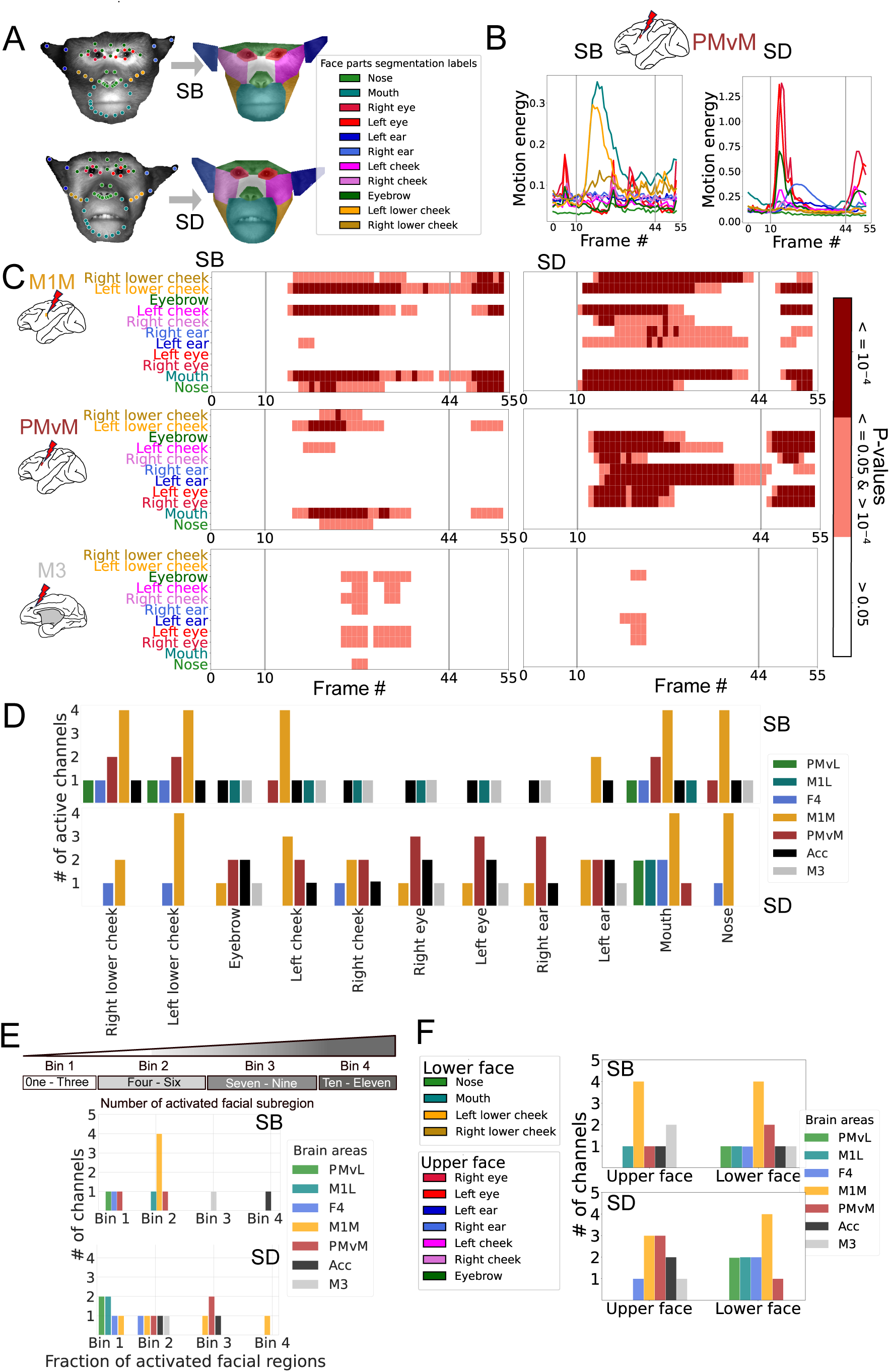
Long-duration stimulation evoked complex movements recruit multiple facial regions. **A.** Video frames with landmarks generated by DeepLabCut (left) and their corresponding segmentation into eleven facial regions. Average motion energy is calculated per segmented region and across trials. **B.** The average motion energy time course for an example stimulation site (PMvM) in both subjects, color-coded by facial subregion. **C.** Significance values for frames from three channels (M1M, PMvM, and M3) across pre-stimulation, stimulation, and post-stimulation periods. Stimulation begins at frame #10 and ends at frame #44, covering the 500 ms stimulation period. Significance values are derived from comparing the no-stimulation data distribution in each facial region to the stimulation data per area (1-tailed t-test, FDR corrected). **D.** Number of statistically significant channels per array that evoked observable movement across the eleven facial regions. **E**. Illustration of four levels of complexity relative to the number of activated facial regions in the top panel. The lower panel shows the number of stimulation channels associated with different levels of complexity for each array, separately for both subjects. **F.** Number of effective channels for each array, evoking activation in the lower and upper face regions separately for both subjects.

To identify statistically significant responses in facial sub-regions, we performed one-tailed t-tests comparing stimulation to “without stimulation” epochs for each facial sub-region and frame. Fig. 3C illustrates examples of this analysis for three cortical sites: M1M (Monkey B: 500 ms, 75 µA; Monkey D: 500 ms, 50 µA), PMvM (Monkey B: 500 ms, 85 µA; Monkey D: 500 ms, 250 µA), and M3 (Monkey B: 500 ms, 150 µA; Monkey D: 500 ms, 300 µA). Fig. 3D summarizes the results for all stimulation channels, organized by their respective arrays. The channels that demonstrated statistical significance (p<=0.05) in the facial sub-regions during the stimulation period were identified as active channels responsible for inducing movements (Fig. 3D). The results indicate that multiple sites engage selective facial sub-regions.

Next, we defined movement complexity by creating four response classes: Bin 1, Bin 2, Bin 3, and Bin 4 (Fig. 3E), based on the number of activated facial sub-regions. Complexity increases with the number of facial sub-regions activated. Generally, most sites did not reach the highest level of complexity (Bin 4), where all facial sub-regions were activated in both subjects. Only one channel from ACC in Monkey B and one channel from M1M in Monkey D reached the highest level (i.e., Bin 4) of response complexity. Most sites showed complexity levels within Bin 1 or Bin 2, indicating the activation of a local subset of facial regions.

Previous anatomical studies, based on the dense projections of the cortical facial motor areas to the facial nucleus, suggest that these areas control either the upper or lower parts of the face (Morecraft et al., 2001). To determine whether FCMS exert strict control over categorical facial regions or coordinate movements of the whole face as needed in natural facial expressions, we characterized which parts of the face were activated by the stimulation of sites in these areas. To quantify this, we categorized the arrays based on their activation of the upper and lower face (Fig. 3F). Our results suggest that arrays in the medial areas (M3 and ACC) predominantly activated the upper face, with some sites also affecting the nose and mouth in the lower face for Monkey B (Fig. 3F). Conversely, all arrays in the lateral brain regions showed activation in the lower face. Some lateral sites, such as PMvM in Monkey D, had a dominant activation in the upper face. Additionally, PMvL and F4 were primarily associated with activation in the lower face. M1L demonstrated activation in the lower face for Monkey B and in both upper and lower face regions for Monkey D. Thus, we found that lateral and medial brain areas activated both the upper and lower face. Further, in agreement with the anatomical study, our functional approach indicate that lateral areas primarily controlled the lower part of the face, while both the lateral and medial facial motor areas controlled the upper part of the face.

Naturalistic facial expressions involve facial muscles synergies that create noticeable movements in specific facial areas. Although we detected activation in various facial sub-regions, we did not identify any noticeable visual similarities between the stimulation-induced movements and the socio-emotional expressions detailed in the literature. For instance, the evoked movements did not resemble common facial expressions such as lip smacking, threat displays, fear grins, yawning, or open mouth play faces, as reported in these studies (Hinde and Rowell, 1962; Maestripieri, 1997; Partan, 2002).

To more directly discern the “naturalness” of evoked movements, we then compared the activation patterns of several facial sub-regions across three natural facial expressions: fear grin, lip-smack, and threat (Sup Fig. 3). First, our analysis revealed that each facial expression had a differentiating activation pattern across facial sub-regions. Second, most sub-regions are engaged at some point in time during the course of the expressions execution (Sup Fig. 3B). This finding contrasts with the results from stimulation-evoked movements, which only activated a subset of facial sub-regions. Therefore, the stimulation-induced behavior did not exhibit the same complexity as typical, naturalistic facial expressions. Although the stimulation elicited movements of mid-range complexity, they did not resemble the full range of natural facial expressions in terms of temporal characteristics or spatial extent.

### Intracortical microstimulation-evoked neural activity propagates to all cortical facial motor areas

Action maps hypothesis implicitly suggest that specific, ethologically defined, motor gestures are mediated through a locally confined network. To test this hypothesis, we took advantage of the fact that while stimulating through one array, we recorded neural activity from the remaining, non-stimulated arrays. This allowed us to assess the spatial distribution of stimulation-evoked neural activity across various cortical areas. Fig. 4A illustrates examples of trial-averaged and z-score multi-unit activity (MUA) recorded from different areas in response to long-train stimulation using one of the channels in PMvM for Monkey B (500 ms, 85 µA) and Monkey D (500 ms, 175 µA). In Monkey D, long train stimulation induced a strong excitation characterized by a rapid onset followed by a gradual decline and return to baseline after stimulation ended. In Monkey B, the stimulation-triggered MUA exhibited changes neural activity in several areas.

**Figure 4.**
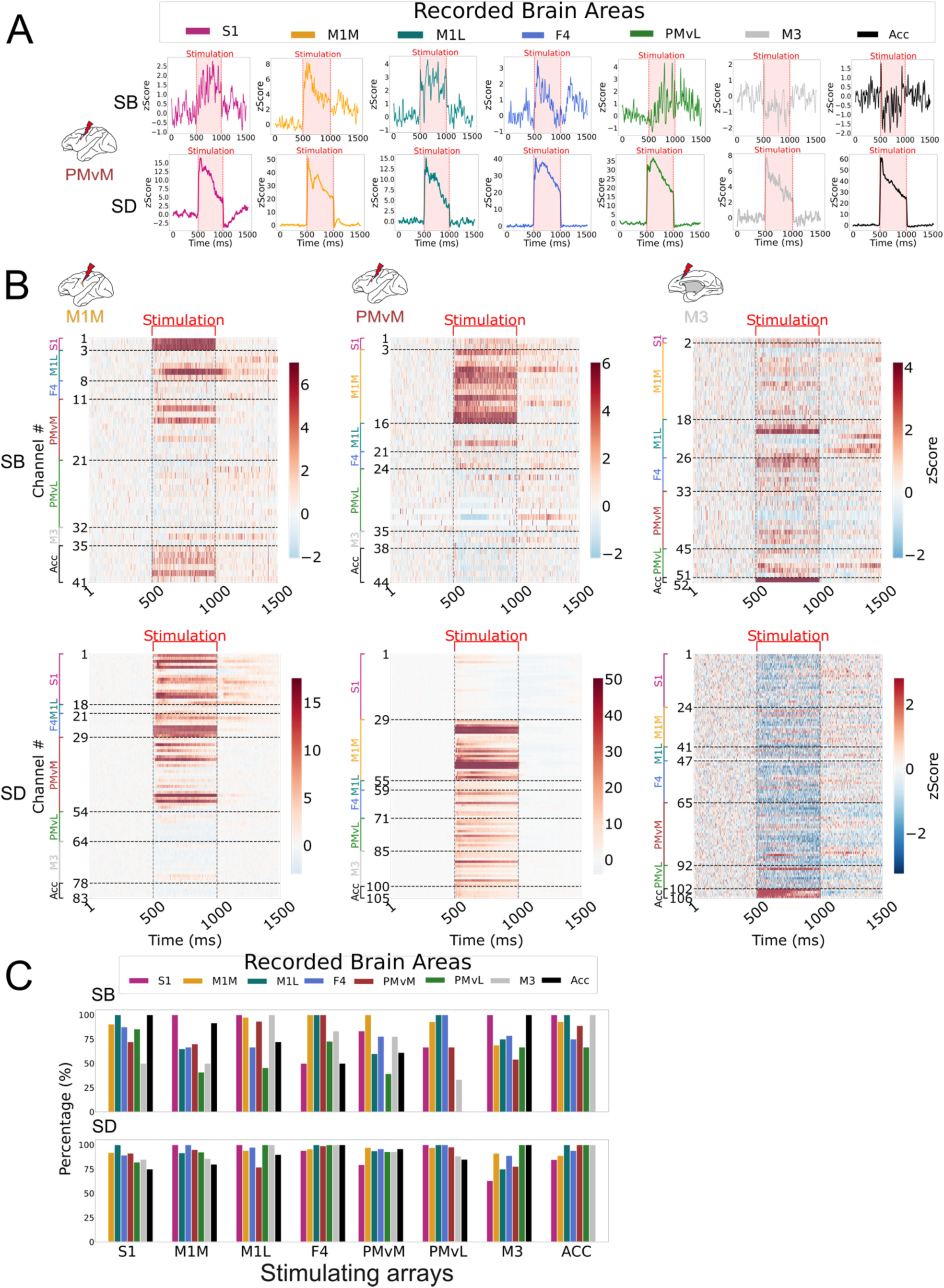
Stimulation-evoked neural responses across different recorded brain areas. **A.** The trial-averaged, z-scored MUA was convolved over time with a Gaussian filter using five standard deviations. Examples shown from channels S1, M1M, M1L, F4, PMvL, M3 and ACC while stimulation was applied on channel 4 in PMvM for both monkeys. Red shading box indicates the stimulation period of 500 ms. **B.** Population peri-stimulus time histograms (averaged, z-scored) shown for two monkeys while stimulation is applied from one of the channels in M1M, PMvM, and M3. The top panel is for Monkey B and the bottom panel is for Monkey D. **C.** Percentage of significantly modulated channels (p<=0.05) out of total active channels driven by stimulation across arrays. Each array had four stimulating channels, and ACC had 2 stimulating channels.

We further analyzed the population responses (trial-averaged and z-scored) at all remaining recording electrodes. Fig. 4B displays the temporal profile of evoked activity while stimulating in electrodes in M1M (Monkey B: 500 ms, 75 µA; Monkey D: 500 ms, 50 µA), PMvM (Monkey B: 500 ms, 85 µA; Monkey D: 500 ms, 175 µA), and M3 (Monkey B: 500 ms, 150 µA; Monkey D: 500 ms, 300 µA). These results indicate that single site stimulation drove time-locked neural activity across various arrays distributed on the cortical sheet. We conducted a 2-tailed t-test (p<=0.05) to determine the statistical significance of changes in responses during the 500 ms stimulation period compared to the 500 ms pre-stimulus period. The results reveal that multiple channels across all recording arrays exhibited statistically significant (p<=0.05) changes in neural activity due to electrical stimulation (Fig. 4C).

Finally, for each subject, we calculated the percentage of significant channels (p<=0.05) relative to the total number of active channels per recording array (Fig. 4C). The results suggest that the effects of stimulation were not confined to nearby sites; rather, stimulation influenced neural activity at sites separated by considerable distances, possibly linked by long-range connections. For instance, stimulation in the medial areas, the ACC and M3, led to significant changes in neural activity across lateral arrays, S1, PMvM, PMvL, F4, M1M, and M1L (Fig. 4C). Conversely, stimulation in the lateral arrays induced notable changes in neural activity in M3 and ACC (Fig. 4C). This indicates that neural activity propagates and induces changes across an integrated network of FCMS. Additionally, we observed modulation of neural activity across the FCMS at the duration of 50 ms (Sup Fig. 4), reinforcing the interpretation that these areas cascade a wealth of shared information, which may be critical for many types of motor control computations.

## Discussion

We examined the organization of stereotypical socio-communicative facial expressions in the cortical facial motor areas using electrical stimulation with two parameters: long-duration microstimulation and supra-threshold current. Such parameters were reported to evoke complex, naturalistic behaviors when delivered to the motor cortex in past studies (Graziano et al., 2002; 2016). Thus, we aimed to determine whether these stimulation parameters could reveal a general ethological organizational principle within the full extent of the facial motor areas. Our findings showed that while these parameters did induce complex facial movements by engaging multiple facial regions at several sites, they did not specifically elicit stereotypical socio-communicative facial expressions. The effects of long-duration stimulation appeared to be an amplified version—both spatially and temporally—of those produced by short-duration stimulation. We found that the evoked neural activity extended across all sampled facial motor areas, suggesting that these areas might function as part of a broad network responsible for generating coherent, context-specific, and socially relevant motor outputs. Our observations suggest that while a single cortical facial motor area may contribute to controlling complex facial movements, it does not seem to be organized into separate zones specifically dedicated to socio-communicative facial expressions.

Our results indicate that short-duration microstimulation elicits simple, localized movements, whereas extending the stimulation duration to 500 ms results in a broader spatial activation pattern that builds on the localized activation observed with 50 ms stimulation. Previous studies have noted that responses to short-duration stimulation are often a truncated version of the more complex movements induced by long-duration stimulation (Stanford et al., 1996; Graziano et al., 2002). This suggests that the complex movements observed with long-duration stimulation might be underrepresented in studies that use only short-duration stimulation. One possible explanation for such discrepancy is that long-duration microstimulation enhances the recruitment of horizontally connected neurons over time, leading to more widespread activation. Indeed, the motor cortex features extensive horizontal connections, which facilitate the spread of activity and the gradual recruitment of various muscle groups (Huntley and Jones, 1991; Keller, 1993; Lund et al., 1993; Capaday et al., 1998; 2009; 2011; 2013). Consequently, the increased facial activation observed during long-duration stimulation may result from the progressive recruitment of additional cortical areas that are linked by horizontal connections around the stimulation site.

Since long-duration stimulation resulted in broader spatial recruitment of facial muscles compared to short-duration stimulation, we further investigated the complexity of the evoked movements related to long-duration condition by assessing the number of recruited facial sub-regions. Our results indicate that the sampled sites recruited various facial sub-regions, encompassing either the upper, lower, or both parts of the face. One organizing pattern revealed by our analysis is that the lower facial regions were mainly controlled by the lateral motor areas, while the upper regions were controlled by both lateral and medial facial motor areas. This result is in line with previous anatomical study which suggested functional division between the lateral and motor areas (Morecraft et al., 2001). Moreover, many sites recruited multiple facial sub-regions, suggesting that cortical facial motor areas do not control local groups of muscles exclusively but instead coordinate several distributed motoneuron pools. Despite this, most of the sites only recruited a sub-set of facial sub-regions and not the whole face, therefore we compared them to the naturalistic socio-communicative facial expressions ethologically described in both their natural habitat and laboratory settings to characterize their degree of similarity (Hinde and Rowell, 1962; Gothard et al., 2004; Morozov et al.,2021). Previous research indicates that naturalistic, socially expressive facial expressions in macaques involve the movement of multiple bilateral upper and lower facial regions (Van Hooff, 1967; Gothard et al., 2004; Mosher et al., 2011). Although the stimulation-induced movements engaged multiple facial sub-regions, they did not closely resemble the socially communicative facial expressions documented in macaques. For instance, we did not observe the rhythmic lip puckering characteristic of lip-smacking or the bilateral open mouth seen in yawning or threatening expressions. Therefore, none of the stimulation sites produced the stereotypical facial expressions used by macaques in social communication. Additional analysis of three natural facial expression in our study also showed that these stereotypical expressions are associated with high-level complexity and recruited most facial sub-regions along the evolution of the required facial movements. In contrast, the stimulation-evoked movements only recruited a more constrained subset of facial sub-regions. These findings suggest that none of the sites explored in this study elicited naturalistic stereotypical facial expressions.

While our experimental approach and results did not identify stimulation-evoked stereotypical facial expressions in the tested sites, future work that includes coordinated stimulation protocols may offer further insights. A plausible interpretation is that the coordination of motor control for naturalistic facial expressions may rely on the dynamic activity of multiple cortical facial areas working together as an integrated neural circuit. Then it follows that if a group of areas functions as a distributed but integrated circuit, we must be able to quantify the functional connectivity between them. To investigate whether the areas sampled in this study are functionally connected, we recorded electrophysiological responses while stimulating each site individually. This approach allowed us to map patterns of functional connectivity between different brain areas. Our results revealed that, regardless of behavior, a coordinated spatial pattern of activity spreads across all sampled areas, although with different temporal activation patterns, suggesting the existence of an internal structure or functional “blueprint” for information processing. Moreover, we also observed the spread of activity regardless of the stimulation duration, as 50 ms trains also caused modulation of activities across different sites. These conclusions held for both subjects, the only difference we noticed was a stronger effect of stimulation in Monkey D compared to Monkey B because we recorded from Monkey D earlier after implantation, which allowed us to collect more neurons and trials per channel. Despite the differences, results from both subjects show a modulation in all recorded arrays. This suggests reciprocal communication between these areas and highlights the need for further investigation into their collective functions to generate naturalistic facial expressions.

A key question is why stereotypical facial expressions did not emerge despite the spread of stimulation-induced neural activity? We observed that stimulation-induced neural activity consistently dominated over baseline activity, often reaching near-saturation tonic levels. While electrical stimulation is effective for altering brain states, it may fall short of replicating the temporally modulated spike trains and specific neural states needed for natural behaviors. Additionally, the computational mechanism described by the balanced neural network model of the motor cortex (Capaday and Van Vreeswijk, 2006; Capaday et al., 2013; Capaday and Van Vreeswijk, 2015) might offer an explanation. According to this model, two populations of reciprocally connected excitatory and inhibitory neurons maintain network dynamics that can quickly adapt to input changes, thereby influencing spike rates (Van Vreeswijk and Sompolinsky, 1996). This model suggests that even if two areas are connected, stimulation of one site might not produce motor outputs from its connections due to local inhibition counteracting local excitation. Moreover, it may be possible to connect different cortical facial motor areas by stimulating them simultaneously. Previous studies, such as those examining simultaneous stimulation in the forelimb representation of cats, showed that the outputs and displacement vectors summed linearly (Ethier et al., 2006). Future research could explore whether simultaneous activation of cortical facial motor areas yields combined movement patterns that more closely resemble stereotypical facial expressions.

Finally, another potential reason we did not observe socio-communicative facial expressions could be related to recent re-evaluations of the traditional topographic models of the motor cortex. Recent studies, such as Gordon et al., (2023), have identified distinct anatomical regions in the motor cortex for representing body movements in more integrative manner versus specific muscle groups. Using fMRI, this research found that the classical homunculus (effector zones) is interspersed with regions that represent task-related movements involving multiple body parts (inter-effector zones). It is possible that the stimulation sites in our study were in the effector regions of the face, rather than in the intereffector regions. This misalignment with the ethological action map hypothesis could be a factor in not eliciting the stereotypical facial expressions, highlighting the importance of accurate localization for producing natural behaviors.

In conclusion, this study represents the first systematic investigation into the representation of stereotypical facial expressions within the facial motor areas. We employed two key criteria—long-duration and suprathreshold electrical stimulation—that were successful in previous studies for eliciting complex naturalistic behaviors. Despite meeting these criteria, we did not observe stereotypical facial expressions. It appears that additional factors may be necessary to effectively stimulate these expressions. Firstly, simultaneous stimulation of multiple cortical facial motor areas might be required. Stimulation of just one area may only activate a limited set of motor synergies, while the combined activity of multiple areas could be necessary to elicit the full range of movement for a facial expression. Secondly, like findings in humans, the macaque cortex may also be organized into effector and intereffector zones. Effective stimulation might need to target the intereffector zones to trigger naturalistic facial expressions. Future studies should explore simultaneous the stimulation of multiple areas and investigate whether the organizational principle of effector and intereffector zones also applies to non-human primates.

## Acknowledgments

Research reported in this publication was supported by the National Institute Of Neurological Disorders And Stroke of the National Institutes of Health under Award Number R01NS110901. The content is solely the responsibility of the authors and does not necessarily represent the official views of the National Institutes of Health. Thanks to W. Zarco for insightful discussion, providing stimulus for natural facial expression and comments on the manuscript. Y. Vazquez, G. Ianni and A. Gonzalez for animal training, surgery planning and preparation. S. Schaffelhofer for respective setups. M.H. Schieber and A.G. Rouse for array implantation surgeries. W. Zarco and W.A. Freiwald for surgeries help. L. Yin for administrative support and the veterinary team of the Rockefeller University for the care of the subjects.

## Declaration of Interests

The authors declare no competing interests.

## EXPERIMENTAL MODEL AND SUBJECT DETAILS

### Subjects

Data was obtained from two adult male long-tailed macaques (Macaca fascicularis), Monkey B (SB) and Monkey D (SD). The procedures followed federal and state regulations and the NIH Guide for Care and Use of Laboratory Animals. Experiments were performed with the approval of the institutional Animal case and Use committees of The Rockefeller University and Weill Cornell Medical college.

## METHOD DETAILS

### Surgical procedures

In a first surgical procedure, standard anesthetic, aseptic and postoperative treatments were used to implant MRI-compatible ceramic screws, acrylic cement and Ultem head post to restrain the head. In a separate surgery, floating microelectrode of 4mm x1.8mm dimensions with 36 platinum/iridium electrodes, or 2.5mm x 1.95 mm dimensions with 18 or 36 electrodes (dense array) were implanted. Among the 36 electrodes in the arrays, 2 electrodes were ground and 2 were reference electrodes. In the case of 18 channel arrays, 1 electrode was ground, and 1 was reference. 32 or 16 channels were used for electrophysiological recordings, and among them 4 or 2 low-impedance (range between 0.010-0.020Mohm) pure iridium electrodes were used for the electrical stimulation. Separation between electrodes was 0.4mm or 0.25mm (only in case of the F4’s array in SD).

The implanted areas are somatosensory area (S1, one 32 active channels array), two arrays in the primary motor cortex medial and lateral halves (M1M and M1L, two 32 active channels arrays), area F4 (F4, one 32 active channels array), two arrays in the premotor cortex ventral medial and lateral halves (PMvM and PMvL, two 32 active channels arrays), one array in rostral cingulate cortex (M3, one 32 active channels) and one array in anterior cingulate cortex (ACC, one 16 active channels array). The procedure followed federal and state regulation and the NIH Guide for Care and Use of Laboratory Animals. At the Rockefeller University, surgeries, electrical stimulation, and electrophysiology were conducted, while functional magnetic resonance imaging was performed at the Citigroup Biomedical Imaging Center (CBIC) of Weill Cornell Medicine. Rockefeller University and Weill Cornell Medicine’s Institutional Animal Care and Use Committees (IACUC) approved all experiments.

### fMRI Imaging

We identified the cortical facial motor areas using the facial movement localizers to surgically guide implantation of floating micro-electrode arrays. Two facial movement localizers were utilized: “the production of communicative social signals” through interaction with videos of conspecifics (Shepherd and Freiwald, 2018), and an orofacial movement localizer through the juice drinking paradigm. Each subject underwent imaging in a 3Tesla Prisma (SIEMENS) scanner. In the “production of communicative social signals”, subjects free-viewed visual stimuli on the screen of conspecifics to produce communicative social signals. In the juice drinking paradigm, Monkey B got juice reward for maintaining fixation at the screen in an event related design. For Monkey D, the subject received juice for a 27 second period (manually delivered at an interval of ∼1to 3 seconds) alternating with 27 second period without juice delivery. During these two paradigms, functional volumes were acquired using custom made Mass General Hospital (MGH) coil with eight receiving channels, using a gradient-echo echoplanar EPI sequence (1.2mm^3 isotropic voxels, repetition time [TR]=2.25 s, echo time [TE] =17 ms, flip angle [FA] =79 degrees). To increase the contrast-to-noise ratio, we injected Molday ION (BioPAL) contrast agent through the saphenous vein immediately before each scanning session. High resolution T1-weighted images were acquired with a magnetization prepared rapid gradient echo (MPRAGE) sequence (0.5 mm isotropic voxels, 240 sagittal slices).

To define facial motor areas in the “production of communicative social signals” localizer, a contrast between facial expression, lip-smack and no-movement or nervous chew was used. In the juice drinking paradigm, facial motor areas were identified based on the contrast between oral-facial movements made during juice drinking versus non-juice drinking periods. Eye position was measured at 60 Hz using a commercial eye monitoring system (Iscan) and facial movements were captured using a MR-compatible infrared video camera (MRC). Both social communication facial expressions and juice drinking behavior were identified and scored manually for their presence in the video recording. High resolution anatomical images were used to align the functional contrast maps and visualize the FCMS. A conjunction activation map was created by utilizing common activation areas between the two paradigms. The demonstrated activation in both the paradigms was used to functionally identify the FCMS and this information was used to guide floating micro-electrodes implantation in the right hemisphere by targeting and calculating positional parameters using the software Cortexplore (cortexplore.com).

### Intracortical microstimulation protocols

Intracortical microstimulation was used to map the evoked movements across the FCMS Two monkeys implanted with chronic micro-electrode arrays were microstimulated during an awake procedure, and they were either moving their face or resting. There were 4 stimulation electrodes per array, except for ACC where only 2 stimulation electrodes were used for electrical stimulation. These selective electrodes were ideal for electrical stimulation because they had low impedance and were made of iridium material. We used an IZ2MH current stimulator from Tucker-Davis Technologies (tdt.com) to pass current through those stimulating electrodes. The output voltage in relation to the electrode impedance was measured to monitor the delivered current.

At the start of the experiments, we identified the threshold current value for the generation of movements (twitches) in each electrode. To determine the threshold current values, the stimulation parameters that were used are cathodal first biphasic pulses with pulse duration of 200 μs in each direction with no gap between pulses at a frequency of 200 Hz for a train duration of 50 ms. We started with a current of 10 μA and increased the current with a step size of 10 μA until any movements in three consecutive trials were observed. Once the threshold values were obtained, we used supra-threshold current values (defined as values above the threshold that caused consistent movements as observed by the experimenter) to collect data from 50 ms duration while keeping the rest of the stimulation parameters the same.

The main goal of the experiment was to identify facial movements linked to 500 ms duration and supra-threshold current values, inspired by Graziano’s successful studies in driving complex behavior at longer durations (Graziano et al., 2002; 2005; 2006). We applied supra-threshold current values (defined as values above the threshold that caused consistent movements as observed by the experimenter) and applied cathodal first biphasic pulses with a pulse duration of 200 μs in each direction with no gap between the pulses at a frequency of 200 Hz at 500 ms train duration. The current values used ranged from 45 μA to 900 μA. Between each stimulation train of 500 ms duration, we used inter-trial intervals of 5s+0.5 ms in Monkey D and 10-30 s in Monkey B. We collected multiple trials within the same day and across days.

### Electrophysiological recordings

To understand the effect of electrical stimulation on the physiological responses of neurons across distributed cortical facial motor areas, we performed simultaneous electrophysiological responses. The electrophysiological recording electrodes were made with either platinum-iridium or iridium material. We recorded from seven arrays while stimulating from one electrode per array. The equipment did not permit simultaneous recordings from the stimulating array. The arrays were connected by cable wires to the pedestals embedded in the acrylic implant. The preamplifier (PZ2) was connected to the head stages (ZCA32), from Trucker-Davis Technologies (tdt.com), which were then linked to the pedestals through an Omnetics connector. The data were sampled at 24.4 kHz, digitized and band-passed filtered (between 300 Hz and 7000 Hz). We synchronized stimulating and recording systems to the same clock to determine the timing of stimulation in relation to electrophysiological data.

Upon inspection of the electrophysiological data, the artifacts related to stimulation were clearly visible in the band-passed filtered data at all recording channels. Therefore, an offline procedure was employed to remove stimulation artifacts from the signal. In this procedure samples from a period of 1.18 ms were removed (blanked) from the start of each stimulation biphasic pulses, and we used linear interpolation to replace the missing samples (Heffer and Fallon, 2008). This was followed by rectification of the signal so that the negative signals become positive. The rectified signal was filtered at 200 Hz with a low pass filter to generate envelopes of multi-unit activity (MUA).

### Videography and analysis

A USB 3.0 monochromatic camera (FLIR Blackfly BFS-U3013Y3M-C) operating at 69.98 Hz was used to record the stimulation-evoked movements. The camera was positioned to capture the front view of the monkey. The size parameter was 1024×1024 pixels, and an infra-red LED from ISCAN (iscaninc.com) to illuminate the face and body. We decided to use videography (as opposed to EMG) because it does not mechanistically affect facial movements and allows us to measure the entire face. While EMG has been utilized in past studies to study facial behavior, these studies only recorded a few superficial muscles while the videography allows to measure the movements of the entire face (Cooke et al., 2003; Shepherd et al., 2012).

### Face tracking with DeepLabCut (DLC)

We used DeepLabCut (DLC) for preprocessing and behavioral data analysis of the videography data (Mathis et al., 2018). We created a new fascicularis primate-face model to track the landmarks on the face using DLC. Among these landmarks, three were placed on each of the ears, ten landmarks on the eyebrow, five on each of the eyes, twelve on the nose, twenty-nine on the mouth, three on each of the cheeks and three on the tongue. The model was trained with 550 frames for Monkey B and 500 for SD on Resnet-50 and was validated.

### Alignment of images

We collected data across several days, which increased the chances for variabilities in the camera and monkey’s location. An alignment procedure was used to correct for the response of any differences related to these factors.

For each monkey, all data for videography were aligned to one single frame with a neutral expression by using landmarks obtained from a subset of DLC points. We performed the alignment using a similarity transformation which includes rotation, translation, isotropic scaling, and reflection in MATLAB. After the alignment of a single frame from each day with the neutral facial expression to a common reference frame, we applied the transformation to the rest of the frame from videos collected on the day of the experimental sessions given that animals are head fixed during the session.

### Estimation of the spatial response field with optic flow

To visualize and define the areas of the face that were moving due to microstimulation, we utilized the optic flow. Optic flow measures the pattern of motion of objects in the visual scene between consecutive frames in the video, caused by relative movements between the object and the camera (Warren and Strelow, 1985; Beauchemin and Barron, 1995). During each session, the monkeys were held head-fixed, and the camera was not moved, so the movements could only be caused by facial movements.

We computed dense optic flow using the Farneback method implemented in OpenCV python on video frames which were first aligned to a common reference and downsampled to a spatial resolution of 512×512 pixels. The dense optic flow gives the magnitude and direction of motion for each pixel of the frame. For further analysis, the magnitude of the optic flow was utilized as the motion energy. Optic flow was calculated throughout a video’s length, which consisted of multiple stimulation trials from the same electrode, interspersed with time without stimulation.

To estimate the spatial response field of the electrical stimulation, we identified the frames corresponding to the stimulation. For each stimulation trial, durations of 500 ms and 50 ms were represented by thirty-four and four video frames, respectively. Next, for each stimulation site, we calculated the mean motion energy across trials for each pixel in the frame, which resulted in the stimulation time response average map. After this step, we recomputed the average using randomly selected trials from periods without stimulation (100 repetitions). The trial average and standard deviation from the “without-stimulation” period were then used to compute the z-score for each pixel of the stimulation time response average. A pixel is considered active when a z-score value reaches two standard deviations for three consecutive frames. To further reduce spatial noise in the activation pattern, a pixel was not considered active if it was not connected by eighty activated pixels in the neighborhood. We used a maximum projection of the z-score over the stimulation duration of either 500 ms or 50 ms to create the spatial response field for the whole face.

We counted the number of activated pixels in the spatial response field and then compared the number of activated pixels during the entire 500 ms and 50 ms durations using a 2-tailed t-test for each subject separately. To better understand whether the spatial activation pattern seen in 50 ms is related to the activation pattern in 500ms, we calculated the overlapping pixels between 50 ms and 500 ms as a percentage of activated pixels at 50 ms.

### Dynamic segmentation of the face regions to identify complexity associated with stimulation evoked movements

The aim of the analysis is to determine the complexity of movements associated with different stimulation sites by studying the effects of long-duration stimulation on different facial sub-regions.

We used a subset of DLC landmarks to establish boundaries for eleven sub-regions. The area under the boundary was defined as a sub-region of the face. These sub-regions were 1) mouth 2) nose 3) right ear 4) left ear 5) right eye 6) left eye 7) right cheek 8) left cheek 9) lower right cheek 10) lower left cheek 11) eyebrow. By using this method, one can follow the contours of the facial sub-regions as they change dynamically during movements.

For each stimulation site, we calculated the average motion energy per facial sub-region at each frame. This procedure was repeated for all trials to obtain a distribution of average motion energy for each facial sub-region across the corresponding frames of the prestimulation, stimulation, and poststimulation periods. Next, I created a null distribution by randomly selecting 100 trials from periods without stimulation and averaging the data across these trials. I repeated this procedure for the no-stimulation period (N = number of trials during stimulation) and averaged the data within each facial sub-region to create a distribution of “without stimulation” for each facial segmentation.

To examine statistical difference between the motion energy of two experimental conditions, “with stimulation” and “without stimulation”, in each of the eleven segmented facial sub-regions, we performed a significant test using the 1-tailed t-test on each frame (time segment). We then used a false discovery rate (FDR), to correct for multiple comparison test for the number of facial sub-regions tested at each frame. A facial sub-region was considered responsive when they showed statistically significant higher motion energy because of stimulation for three consecutive frames.

To identify the complexity of movements associated within each stimulation site, we first operationally defined the complexity of movements by the number of activated facial sub-regions. We created a four-level classification of complexity: Bin 1, Bin 2, Bin 3 and Bin 4. Bin 1 contained activation of one to three facial sub-regions. Bin 2 contained activation of four to six facial sub-regions, Bin 3 contained activation of seven to nine facial-sub regions, and Bin 4 contained activation of ten to eleven facial sub regions. For each stimulation site, we counted how many facial sub-regions were activated and assigned the results to their respective bin.

Next, we investigated whether the arrays activated the upper or lower parts of the face by analyzing the distribution of active channels per array for these two parts of the face. The upper part of the face includes the right eye, right ear, right cheek, eyebrow, left eye, left ear, left cheek. The lower part of the face includes the mouth, nose, left lower cheek, and right lower cheek. If a channel activated even one of these upper or lower face parts, we considered it responsive for upper or lower facial movements.

### Dynamic segmentation of the face regions to identify complexity associated with natural facial expressions

The aim of the analysis is to compare the complexity of naturalistic facial expression with complexity of stimulation evoked movements. We analyzed four facial expressions from a single monkey (separate from the two-monkeys used for electrical microstimulation experiments). These facial expressions were fear grin, lipsmack, threat and neutral. We used DLC landmarks to establish boundaries for eleven sub-regions, and calculated motion energy per region. The videos for each facial expression had 120 frames. For each frame, we computed the z-score of motion energy for these facial expression in each sub-region, based on the mean and standard deviation of the motion energy from the neutral expression. A facial sub-region was considered recruited only if the z-score exceeded 2 standard deviations for 3 consecutive frames. If any facial sub-region was active at a point during the facial expression, then we considered it as a region activated by the facial expression.

### Dimensionality reduction of behavior

The next question we aimed to answer was whether evoked movements associated with 50 ms and first 50 ms period of 500 ms (early epoch) are similar? We decided to project the higher dimensional data into a lower dimensional space to simplify the visualization of temporal and spatial dynamics of stimulation evoked movements. We used the Potential of Heat-diffusion for Affinity-based Trajectory Embedding (Phate) to apply dimensionality reduction to the trial averaged and z-scored motion energy of stimulation-evoked video frames. Phate creates a nonlinear embedding of the high-dimensional data while highlighting the continual nature of the underlying trajectories (Moon et al., 2019). The major advantage of Phate is that it represents both the local and global structure of the data while embedding it into lower dimensions. In contrast, another popular method like Principal component analysis (PCA) is limited to the global structure of data, meaning local information can be lost because of noise and non-linearity of the data structure (Moon et al., 2019). t-Distributed stochastic neighbor embedding (TSNE) only considers the local structure of the data and collapses the trajectories into clusters (Moon et al., 2019). Compared to PCA and TSNE, Phate algorithm allows linear or non-linear clusters, branches, and progressions to become clear in a lower-dimensional space (Moon et al., 2019). Thus, Phate enables the reduction of high-dimensional pixel space into a more manageable latent space, making it easier to compare trajectories by minimizing redundancies and capturing the essential structure of the data.

To perform the analysis, we first extracted the pixels per frame containing the trial-averaged and z-scored motion energy (z-score calculated using the mean and standard deviation of randomly selected trials, repeated 100 times, during the “without-stimulation” period) from the stimulation periods of 500 ms and 50 ms. These pixels were then arranged into a vector form, referred to as M_OF_. Subsequently, we constructed a matrix consisting of thirty-four vectors, which corresponded to the product of the number of pixels in each frame and the number of frames during the stimulation period (M_OF x_ N_frame_). We repeated the same procedure for 50 ms duration consisting of four vectors corresponding to the stimulation period and created the matrix (M_OF x_ N_frame_). Next, we concatenated the matrices of 500 ms and 50 ms together to obtained a (M_OF_ x N_frame)_ x N_video_. N_frame_ corresponds to the number of frames in the stimulation time, and N_video_ corresponds to the number of stimulation durations (500 ms and 50 ms). To facilitate a comparison of the movement trajectories between the 500 ms and 50 ms durations at the same stimulation site, we projected the data for both durations into a common Phate space. This analysis was repeated for the sites that produced responses in both the 500 ms and 50 ms durations.

Following, we examined whether the evoked movements at 50 ms is a truncated version of the response at 500 ms. To begin with, we concatenated the 3-dimensional trajectory from the Phate space for 50 ms into a one-dimensional vector, and another one-dimensional vector for the first 50 ms of the 500 ms (early epoch). We performed Spearman rank correlation between the vectors of 50 ms duration and the first 50 ms duration of the 500 ms to evaluate their similarities. To determine the statistical probability of our observations, we conducted a permutation test by comparing the observed test statistics to the distribution of test statistics generated from permuting the data 100,000 times. Statistically significant correlation values suggest that the movement associated with the 50 ms is a shortened version of the 500 ms duration.

### Concurrent neural response measurements

The aim of the analysis is to evaluate the spread of evoked neural activity across simultaneously recorded channels in seven different arrays. At the start of each experimental session, we quantified the number of channels which showed spontaneous spikes and classified them as active channels. We considered a channel as active if it passed the threshold of five standard deviations multiplied by root mean square of the signal. The channels that were active were included for the analysis.

The preprocessed continuous MUA data were used for the analysis. The extracted time series was examined in three relevant epochs of 500 ms each: pre-stimulation, stimulation, and post-stimulation activity (a total of 1500 ms). We applied the same approach for the 50ms stimulation duration condition, resulting in a total of 1050ms. We generated a trial averaged MUA activity for each of the recording electrodes per array. The time series normalization (z-score) was performed, by using the mean and standard deviation extracted from the pre-stimulation epoch. Only for visualization of graphs for individual channels at Fig. 4A and Sup Fig. 4A we convolved the signal with a gaussian filter with standard deviation of 5. For the visualization of z-scores in population peri-stimulus histograms, we used python’s Seaborn package to compute the colormap range with robust quantiles instead of the extreme values Fig. 4B and Sup Fig. 4B.

For 500 ms duration, we assessed the MUA’s modulation in two main experimental conditions: stimulation and pre-stimulation. The pre-stimulation condition was defined as the time prior to stimulation comprising 500 ms preceding the electrical stimulation onset (baseline period). We first averaged the values across the epoch of 500 ms time window for the pre-stimulation and stimulation epochs, for each condition, and created a distribution over the trials. Using a 2-tailed t-test (p<=0.05), we tested the significance of the two experimental conditions. We calculated the percentage of significantly modulated channels out of total active channels driven by stimulation across arrays. Following the analysis of neural data related to 500 ms duration, we applied the same approach for the neural responses to 50 ms train duration. The pre-stimulation condition was defined as the time prior to stimulation comprising 50 ms preceding the electrical stimulation onset (baseline period).

---

**Sup Figure 1.**
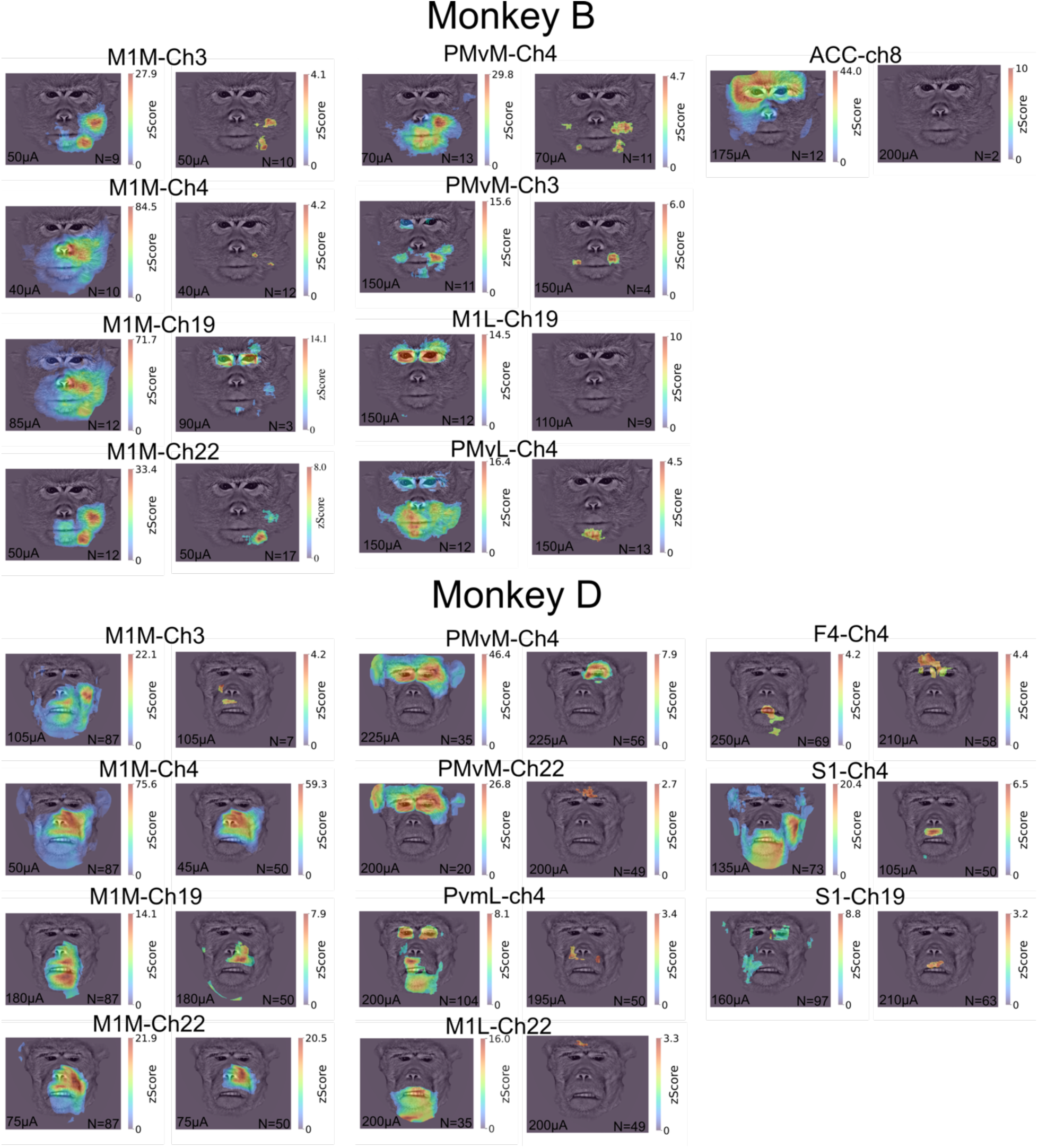
Stimulation-evoked responses overlaid on the face. **A.** The STRA map was computed for each pixel within the face and z-scored by the data from “without-stimulation” epoch. The overlay heat map displays the maximum z-score across the entire stimulation period of 500 ms (Left panel) and 50 ms (Right panel) for each (Monkey B on top, and Monkey D on the bottom)

**Sup Figure 2.**
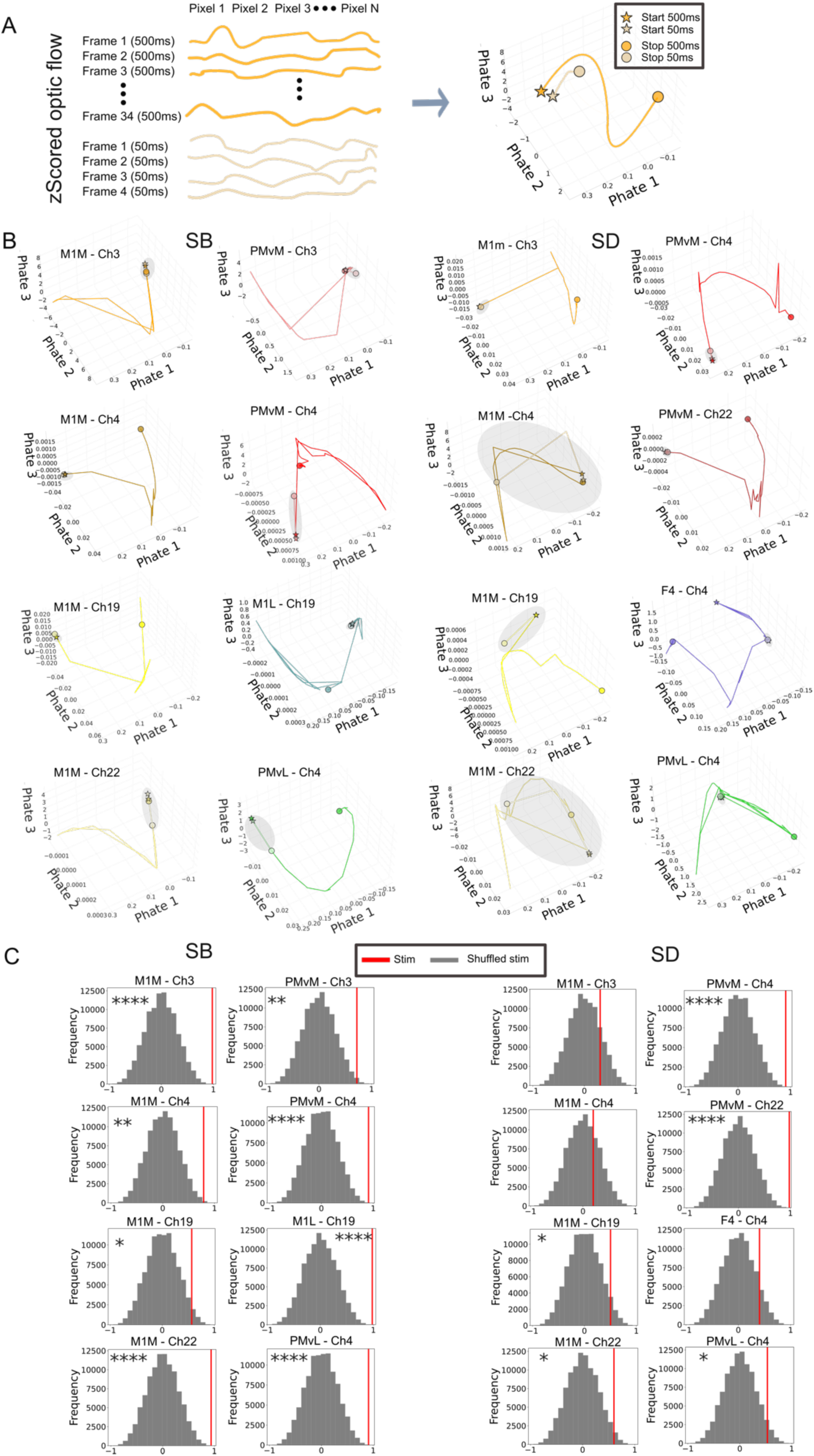
**A** Schematics of the Phate dimensionality reduction analysis on trial-averaged and z-scored motion energy time-series for both durations (500 ms and 50 ms). **B.** Phate dimensionality reduction for stimulation channels. The grey circle is used to visualize the location of the trajectory corresponding to 50 ms. **C.** Permutation null distributions relative to the observed experimental statistic (red vertical line). The trajectories similarity across stimulation conditions was assessed with the Spearman’s rank correlation. (shuffled 100,000 times, *p<0.05, **p<0.01, ***p<0.001, ****p<0.0001). The left panel shows Monkey B and the right panel shows Monkey D.

**Sup Figure 3.**
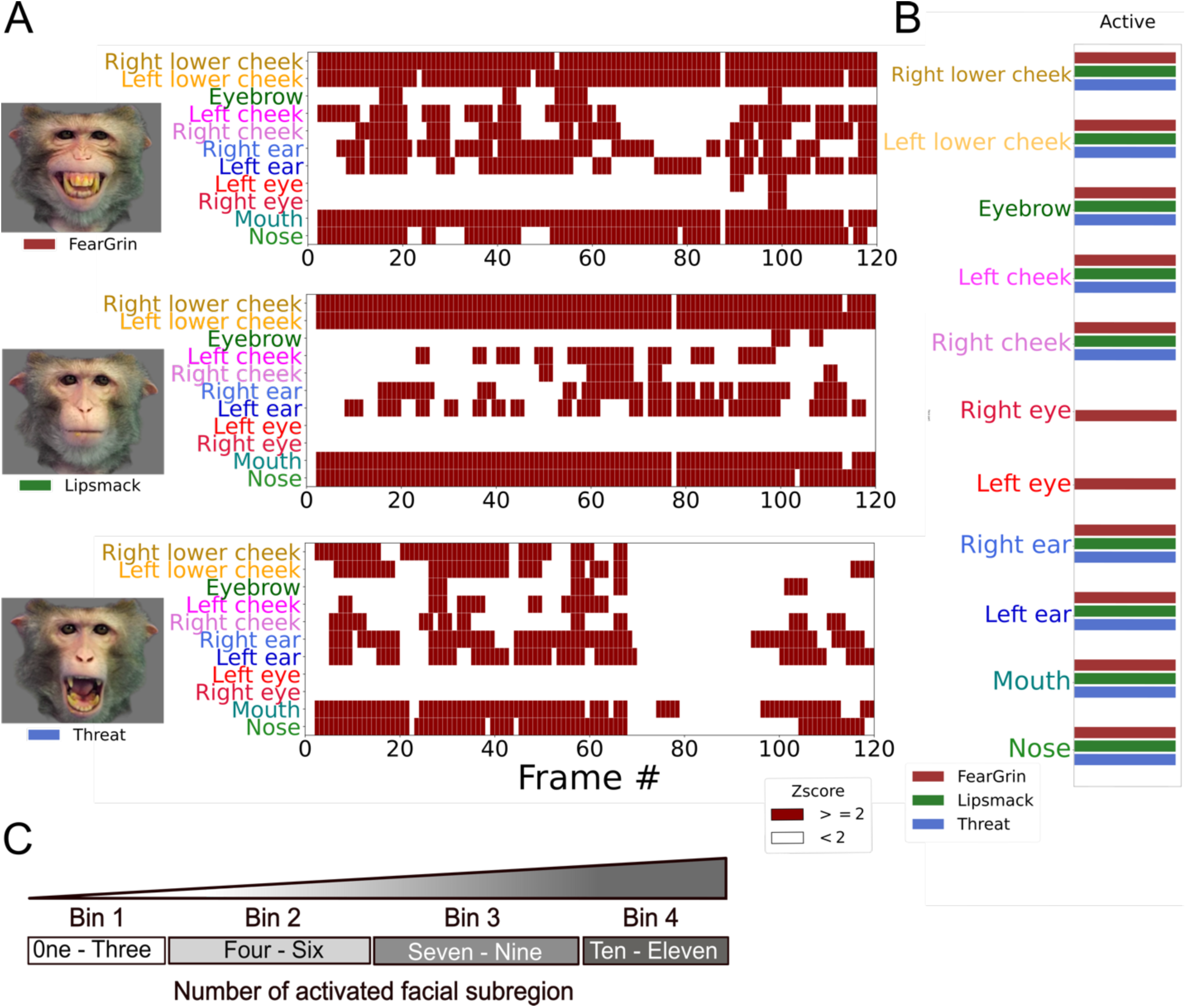
**A.** Activation pattern of facial sub-regions during three naturalistic facial expressions, fear grin, lipsmack and threat from a different subject. Each facial expression has a unique fingerprint of activation. **B.** Facial sub-regions that show activation at one point during the expression’s evolution. **C.** The complexities observed in the three facial expressions were associated with Bins 3 and 4, suggesting that these expressions activated most of the facial sub-regions.

**Sup Figure 4.**
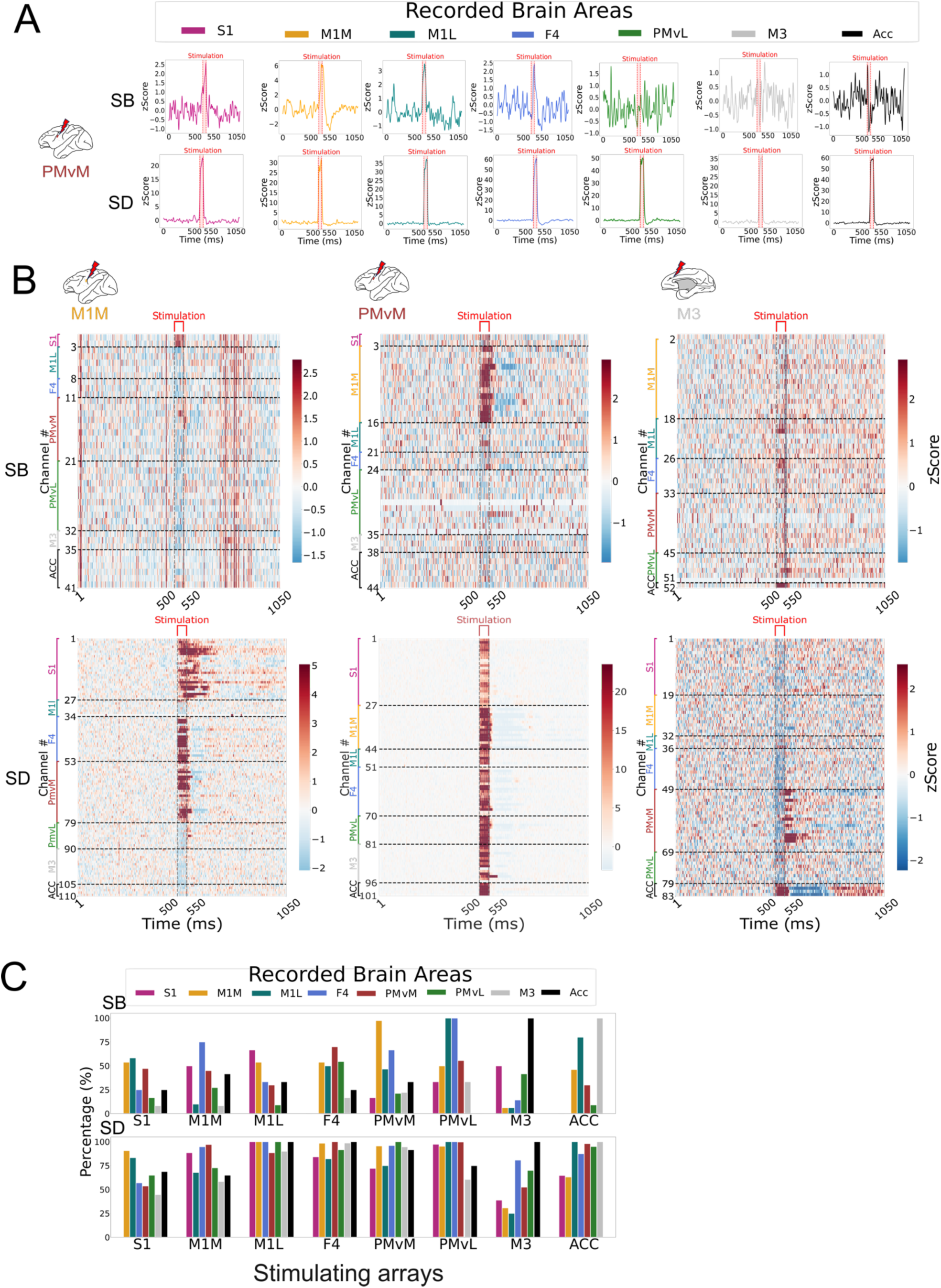
Stimulation-evoked neural responses across different recorded brain areas. **A.** The trial-averaged, z-scored MUA was convolved over time with a Gaussian filter with five standard deviations. Examples shown from channels S1, M1M, M1L, F4, PMvL, M3 and ACC while stimulation was applied on channel 4 in PMvM for both monkeys. Red shading box indicates the stimulation period of 50 ms. **B.** Population peri-stimulus time histograms (averaged, z-scored) shown for two monkeys while stimulation is applied on one of the channels in M1M, PMvM, and M3. The top panel is for Monkey B and the bottom panel is for Monkey D. **C.** Percentage of significantly modulated channels out of total active channels driven by stimulation across arrays. Each array had four stimulating channels, and ACC had 2 stimulating channels.

## Notes

### Competing Interest Statement

The authors have declared no competing interest.

